# On the inference of a southern origin of the North American firefly *Photinus pyralis*

**DOI:** 10.1101/851139

**Authors:** Ana Catalán, Sebastian Höhna, Sarah E. Lower, Pablo Duchen

## Abstract

The firefly *Photinus pyralis* inhabits a wide range of latitudinal and ecological niches, with populations living from temperate to tropical habitats. Its ample geographic distribution makes this species an ideal system for the study of local adaptation and demographic inference of wild populations. Therefore, in this study we modelled and inferred different demographic scenarios for North American populations of *P. pyralis*, collected from Texas to New Jersey. To do this, we used a combination of ABC techniques (for multi-population/colonization analyses), and likelihood inference (*dadi*) for single-population demographic inference, which proved useful with our RAD data.We uncovered that the most ancestral North American population lays in Texas, which further colonized the Central region of the US and more recently the North Eastern coast. Our study confidently rejects a demographic scenario where the North Eastern populations colonized more southern populations until reaching Texas. Our results suggest that *P. pyralis* originated in Central- or South America, followed by migration events that populated northern latitudes. Finally, modelling the demographic history of North American *P. pyralis* serves as a null model of nucleotide diversity patterns, which will inform future studies of adaptation, not only in *P. pyralis*, but also in other North American taxa.

## Introduction

The current distribution of a species is the result of the intricate evolutionary history that often characterizes natural populations. A number of past and present factors, such as geological events (e.g. the formation of land bridges or mountain ranges), ice ages (e.g. the change of ice sheet area across the glove) and more recently, the active migration of populations into novel habitats (Mann 2007; Roy and Lachniet 2010) determine the distribution pattern of a species. In the Americas, the present distribution of flora and fauna was strongly influenced, firstly by the geographic isolation of the land masses of North and South America and secondly, by their subsequent union. The uprise of Central America, a process that started 12 million years ago (MYA) and was completed ∼ 2.7 MYA (Lessios 2008), enabled and accelerated biotic interchange. Additionally, more recent events such as orogeny and glaciations have further defined the distribution of the biotic community in the American continent (Gonzalo 1987).

To understand how past events have determined the distribution of a species and how species distribution is constantly shaped by more recent environmental changes, we need to investigate different taxa. The investigation of different taxa, with distinct generation times, vagility, reproduction and ecological strategies, will widen our understanding into why some species have a very broad or even world-wide distribution, whereas others are endemic being only able to live under specific conditions. In this study, we investigate the demographic history of the firefly *Photinus pyralis*. *P. pyralis* has a wide distribution range, with documented collections from Ontario, Canada (Fallon et al. 2018) to Venezuela in South America (The Bavarian State Collection of Zoology). Thus, *P. pyralis* offers an excellent opportunity to investigate how an insect can acquire such a broad geographic distribution and how the dynamics across populations are evolving.

The wide geographical distribution of *P. pyralis* suggests that this species is highly vagile and adaptable. For example, populations thriving in northern latitudes have to withstand harsh winter conditions. To cope with winter, northern populations of *P. pyralis* overwinter as larvae, a physiological response that is not needed in populations that live in warm areas, such as Texas, and in Tropical regions such as in Central America (Lloyd 1966; Fallon et al. 2018). The ample spectrum of ecological niches and geographical latitudes that *P. pyralis* inhabits makes this species suitable to elucidate the wide range of insect colonization patterns in the Americas as well as adaptation events. In the case of *P. pyralis*, it is not known where the most ancestral population of the species resides, knowledge that would give us information about its place of origin as well as of its dispersal history.

Fireflies are conspicuous beetles that produce bioluminescence as an aposematic signal against predators (Lewis and Cratsley 2008), to attract prey (Vencl and Carlson 1998) and to allure potential mates (Lewis and Cratsley 2008). Because of their production of light, with the exception of some firefly species that have lost their bioluminescence and have become diurnal (Branham and Wenzel 2003), firefly populations are easy to spot, facilitating the study of their biology and behavior. *P. pyralis* lives for one to two years as larvae preying upon snails and earthworms until they metamorphose into adult fireflies during the summer months in Northern latitudes (Faust 2017) and during the rainy season in Tropical habitats (Schuster 1997).

In the present study we investigated the demographic history of *P. pyralis* by analyzing RADseq data from 12 populations collected in North America, which range from Texas to New Jersey. Previous work revealed population structure among these populations as well as heterogeneous levels of nucleotide diversity and genetic distances across populations (Lower et al. 2018). For example, Lower et al. showed that the Texan population has the highest nucleotide diversity and the highest levels of population differentiation when compared to the other populations examined, hinting that the Texan population might be the oldest of North America (Lower et al. 2018). Beside the genetic diversity found across populations, phenotypic traits such as life cycle length (Fallon et al. 2018), flash color (male light emission peak wavelength; Sander and Hall 2015), and body size (Lloyd 1966) have been described or hypothesized as polymorphic traits across *P. pyralis* populations. Whether this variation is due to adaptation, genetic drift and/or demography is still to be elucidated.

By using the previously generated population data and by generating population genetic summary statistics we proposed and tested various demographic scenarios for *P. pyralis*. With this work, we generated the first hypotheses regarding the demographic history of North American populations of *P. pyralis* and hypothesize on the putative most ancestral population of this species. Furthermore, characterizing the nucleotide diversity patterns left by demography alone sets the base to detect nucleotide patterns that deviate from neutral expectations. Having a null expectation of nucleotide diversity will enable us to detect the local adaptation events that permitted *P. pyralis* the colonization of a great variety of natural habitats in future studies.

## Materials and Methods

### Clustering for demographic modelling

The data consists of single nucleotide polymorphisms (SNPs) generated with double digest restriction-site associated (ddRAD) sequencing data from 15 individuals from each of 12 populations of *P. pyralis* from North America (Table 1). A total of 2019 SNPs were kept for downstream analyses after quality control and subsequent filtering steps (Lower et al. 2018). Following the population genetic analysis presented in that study, and current geographical location of the sampled populations, we propose three population categories: Eastern, Central, and Texan. To test for genetic congruence of these three clusters we quantified the probability that the different populations were assigned to one of the clusters by using a Discriminant Analysis of Principal Components (DAPC) (Jombart et al. 2010), as implemented in the R package *adegenet* (v1.4.2) (Jombart and Ahmed 2011). After pooling, the Eastern population consisted of n=66 individuals, the Central population with n=63 individuals, and the Texan population with n=25 individuals, yielding a total of 154 sampled diploid individuals (308 chromosomes).

**Table 1.**
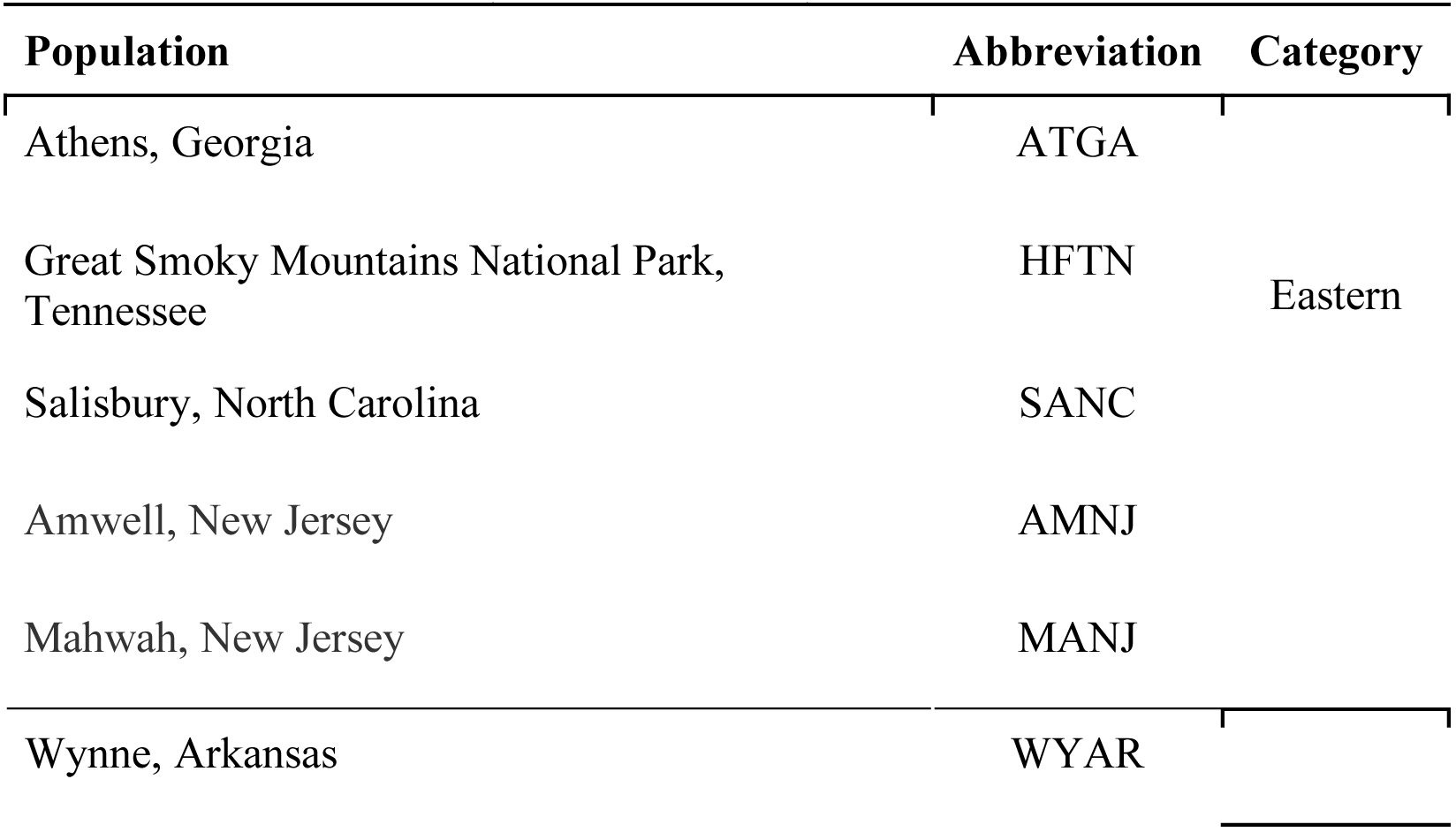
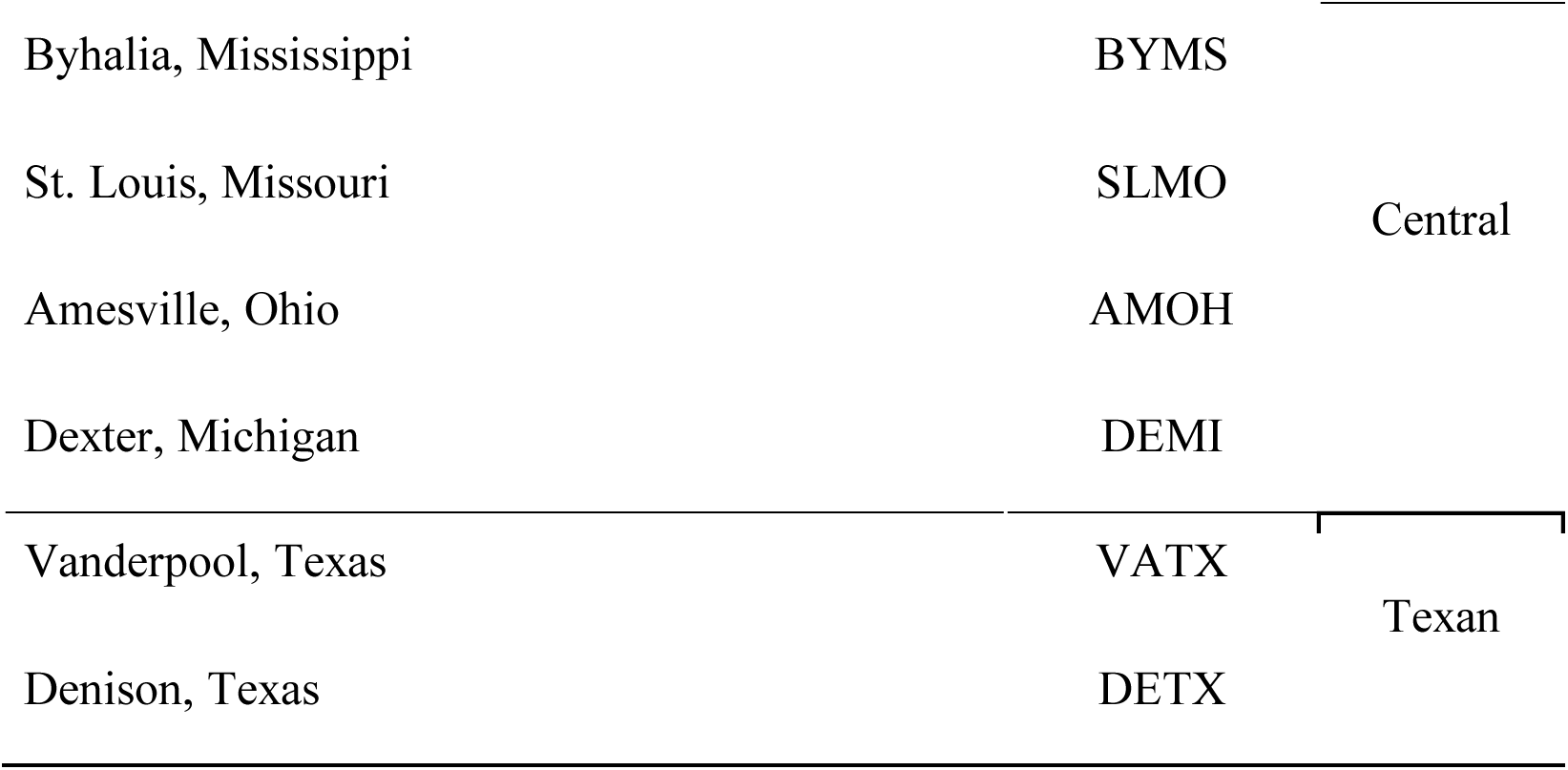
Populations sampled, their abbreviations, and their respective assigned category: <u>Eastern, Central, or Texan (Lower et al. 2018).</u>

### Data preparation for demographic analyses

In our case, it is necessary to take some additional steps with the RAD data in order to make it suitable for a demographic analysis. We are aware that, ideally, a demographic analysis benefits the most from neutral sites, where selection does not confound the effects of demography, and where the maximum number of polarized SNPs are available (the more data, the more power to infer demographic parameters, (Excoffier et al. 2013)). Our data, however, consists of SNPs that are randomly spread throughout the genome, where selection has most likely played a role in maintaining that diversity. Additionally, it is known that next-generation filtering pipelines affect the low-frequency variants more than the rest of sites (Duchen et al. 2013). For these reasons we applied the following procedures to ensure an unbiased demographic analysis with the RAD data at hand: 1) From the original set of 2019 SNPs we discarded all singletons, or class “1” of the site-frequency spectrum (SFS), for downstream analyses. Variant-calling in next-generation sequencing affects the other SFS classes equally, so there is no bias (Duchen et al. 2013). 2) Because there is no outgroup sequence we summarized the data in statistics that are unaffected by polarization (see below). In our case, this implies that excluding singletons means that we also excluded class “n-1” from the SFS. Finally, 3) we performed simulations including selection to account for the evolutionary force that helped maintain many of the variants present in our data (see section “ABC simulations”).

### Observed summary statistics

From the observed data, we calculated the following summary statistics: number of segregating sites *S*, Watterson’s *θ*_W_ (Watterson 1975), *π* and distance of Nei (Nei and Li 1979), Tajima’s *D* (Tajima 1989), linkage disequilibrium *Z*_nS_ (Kelly 1997), the folded SFS, Weir-Cockerham’s *Fst* (Weir and Cockerham 1984), and the Wakeley-Hey “W” summaries of the joint folded SFS (Table 2 and Table 3) (Wakeley and Hey 1997). As explained before all the above-mentioned statistics are unaffected by the polarization of the observed SNPs. The demographic models tested are described in the results section (Figure 2). Here, statistics *S*, *θ*_W_, and *π* summarize nucleotide diversity, Tajima’s *D* and W summarize the SFS, *Z*_nS_ summarizes linkage disequilibrium, and the distance of Nei and *Fst* describe population differentiation.

**Table 2.**
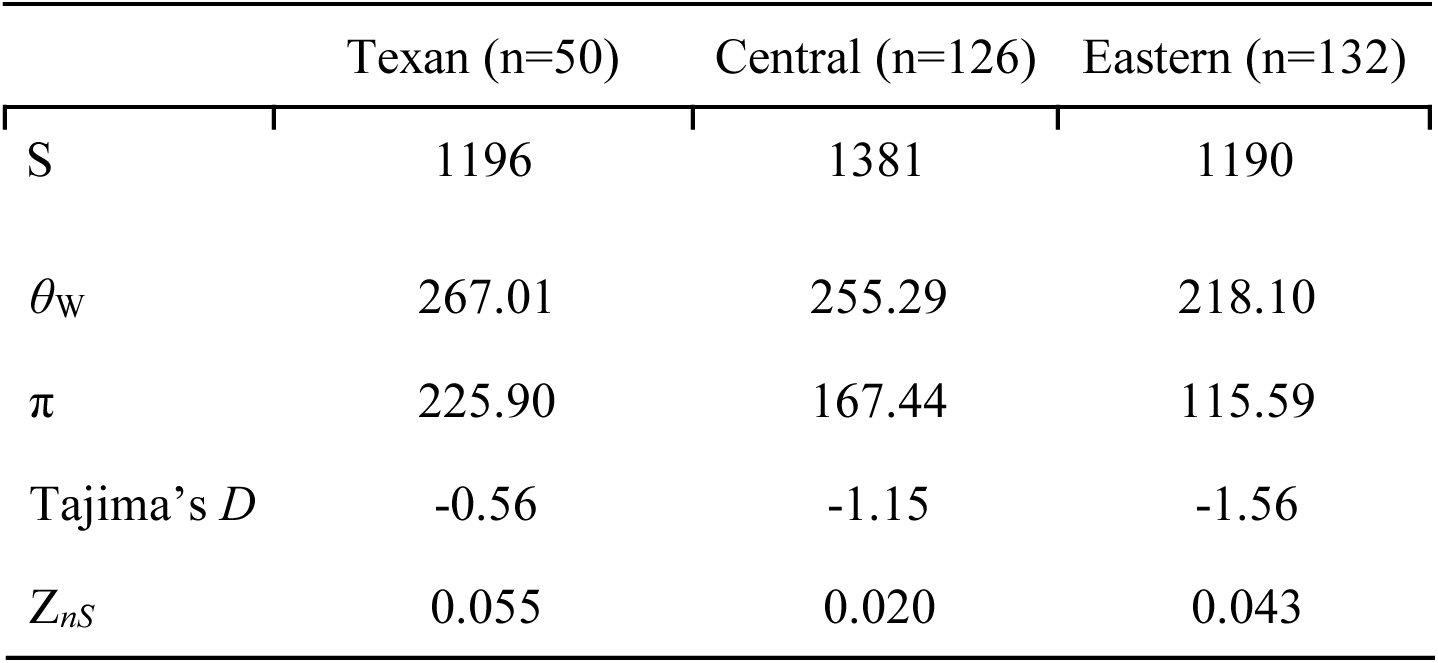
Observed summary statistics for the entire set of loci. Sample sizes (chromosomes) per population are given in parentheses.

**Table 3.**
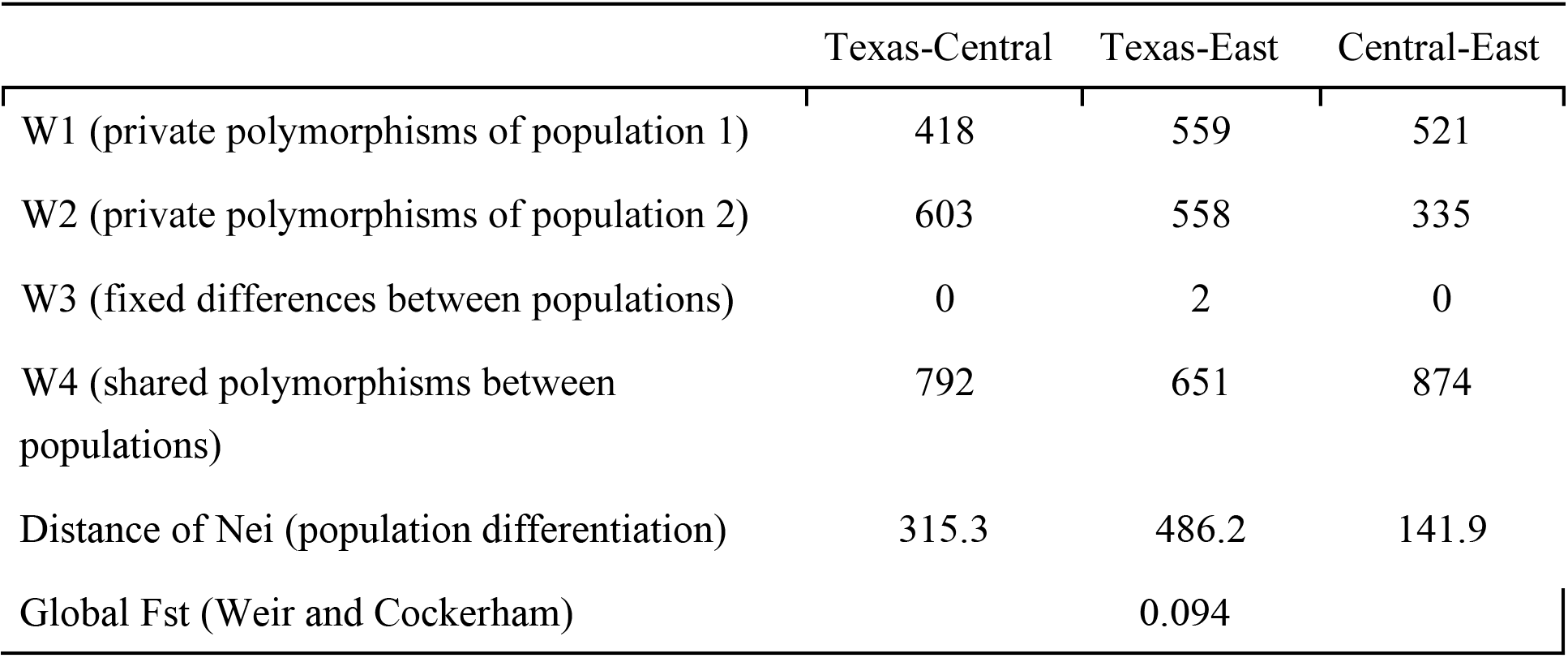
Population pairwise statistics. Wakeley-Hey W statistics summarizing the JSFS, plus population differentiation statistics.

### Simulations

To recreate the variant-generation process of our RAD data we simulated exactly 2019 SNPs (for 308 chromosomes: 50, 126, and 132 chromosomes representing the Texan, Central, and Eastern populations, respectively), for each of the four demographic models described in Figure 2 (see Results for a description of each model). These coalescent simulations were done with the program *msms* (Ewing and Hermisson 2010). To exactly recreate the treatment given to the observed data (see section Data preparation for demographic analyses) from the simulated sites, we removed the SFS classes “1” (singletons) and “n-1” for each population, and calculated all summary statistics described above. Recall that these summary statistics do not depend on the polarization of the data, that is, their values are the same whether we know which alleles are ancestral or which ones are derived. Given that the observed SNPs coming from RAD sequencing are spread throughout the genome (including both conserved and neutral regions) we expect that some degree of selection has helped maintain that diversity for many of those sites. For this reason, we included the *msms* switch-SAa with a value of selection drawn from a prior distribution. This prior distribution, plus the prior distributions for the other parameters are shown in Table 4. We repeated the simulation procedure 50,000 times for each demographic model.

**Table 4.** Parameter estimates and their respective priors. These posterior distributions are shown in Fig. 4. Here, *Ne* stands for effective population size, *T*_CE_ corresponds to the split time between the Central and Eastern populations, *T*_TC_ corresponds to the split time between the Texan and Central populations, and *S*_Aa_ is the selection coefficient.

| Parameter | Prior | Mode | 95% quantiles |
| --- | --- | --- | --- |
| $\log_{10}(Ne_{Central}/Ne_{Texas})$ | unif(-3,6) | 0.155 ($Ne_{Central} = 1.43 * Ne_{Texas}$) | (-0.30,2.78) |
| $\log_{10}(Ne_{East}/Ne_{Texas})$ | unif(-3,6) | -2.702 ($Ne_{East} = 0.0020 * Ne_{Texas}$) | (-3.00,-0.24) |
| $\log_{10}(T_{CE})$ | unif(-3,0) | -2.323 (= $0.0048 * 4Ne_{Texas}$ generations ago) | (-3.00,-1.97) |
| $\log_{10}(T_{TC})$ | unif($\log_{10}(T_{CE}),1$) | -1.591 (= $0.026 * 4Ne_{Texas}$ generations ago) | (-2.62,-0.88) |
| $\log_{10}(S_{Aa})$ | unif(0,4) | 1.239 | (0.26,3.63) |

### Model choice

To estimate confidence in the estimated models, we used Approximate Bayesian computation (ABC), with all 50,000 simulations per model to calculate the posterior probabilities of each of the four demographic scenarios using the R package *abc* (Csilléry et al. 2012). Model choice was based on the following summary statistics: *θ*_W_, *π*, Tajima’s *D*, *Z*_nS_, W statistics, distance of Nei, and Fst (we did not use the number of segregating sites *S* because it is directly correlated with *θ*_W_, or the folded SFS because it is well summarized with the W statistics and Tajima’s *D*). We validated the power of the model-choice procedure by sampling a random vector of “pseudo-observed” summary statistics from the simulations and re-calculating the probability of them coming from one of the four models. We performed this validation 100 times and scored the number of times the right model was chosen. Next, we performed model choice for the observed vector of summary statistics. Finally, to check how well the best model can predict the observed data we plotted the distributions of summary statistics under the best model against the corresponding observed statistics (Figure S2).

### Parameter estimation

Parameter estimation was accomplished by using both the rejection (Tavare et al. 1997; Pritchard et al. 1999) and regression (Beaumont et al. 2002) algorithms using the R package *abc*. Briefly, we kept (“as accepted”) the closest simulations to the observed summary statistics and generated distributions of their associated parameters. These distributions represented an approximation of the posterior probability of each particular parameter based on the “rejection” algorithm (Tavare et al. 1997; Pritchard et al. 1999). Then, by performing a local regression between the accepted simulations and their corresponding parameters an improved version of this posterior probability can be generated (Beaumont et al. 2002). To reduce the high dimensionality of the summary statistics while keeping the maximum amount of information still available we used partial least squares (pls) in an ABC context (Wegmann et al. 2009). The ABC regression step for parameter estimation was performed with the pls-transformed statistics. A validation of parameter estimation was also performed by using the pseudo-observed statistics from the simulations and re-estimating the parameters with the regression algorithm.

### Demographic analysis for single populations

With ABC we focused on multiple-population colonization scenarios (Figure 2). However, to further investigate the demography of each single population (and to unburden the amount of parameters of the multiple-population models) we also analyzed our data using *dadi* (Gutenkunst et al. 2009). To be more precise, we fitted five different, commonly used single-population models: (1) Neutral equilibrium model; this is the null-model and assumes a constant population size. (2) Two-epoch model; this model describes one population-size change that could either be a decrease or an increase. (3) Growth model; this one describes an exponential population-size change reflecting exponential growth or decline. (4) Bottle-growth model; this model describes a bottleneck (or expansion) event followed by a population growth/decline. (5) Three-epoch model, which describes two population-size changes in time. As for our previous analyses, we used the option in *dadi* to apply the folded SFS since we cannot polarize the SNPs, and masked out singletons. For each model we calculated the maximum likelihood parameter estimates and computed the optimized likelihood. This likelihood is used to select the best fitting model, for example, applying the Akaike Information Criterion (AIC).

### Data retrieval and processing for node calibration

To retrieve sequences of fireflies we did a GenBank search using the taxon tag “Lampyridae” (fireflies). Sequences belonging to this taxonomic group were clustered in putative orthoclusters using the UCLUST algorithm (version 6.0.0) (Edgar 2010) as implemented in SUMAC (version 2.2.0) (Freyman 2015). Orthoclusters were aligned with MAFFT (version 6.2.40) (Katoh et al. 2002) and badly aligned regions were removed with Gblocks (version 0.91b) (Talavera and Castresana 2007). Sequence duplicates were removed and further alignment curation was done manually with AliView (version 1.18.1) (Larsson 2014). After curation, five orthoclusters representing five genes were chosen for phylogenetic node calibration: *Cytochrome c oxidase subunit I* (COI, length:1478nt, sequences:123), *28S ribosomal RNA* (28S, length:632, sequences:28), *16S ribosomal RNA* (16S, length:473, sequences: 100), *wingless* (WG, length:404, sequences:55) and *carbamoyl-phosphate synthetase 2* (CAD, length:1763, sequences:41). These genes have been used to diagnose species-level relationships in fireflies (Stanger-Hall et al. 2007; Sander and Hall 2015; Stanger-Hall and Lloyd 2015; Martin et al. 2017).

### Calibrating the node to P. pyralis

To increase our understanding on the population history of *P. pyralis*, we estimated a time-calibrated phylogenetic tree using the software RevBayes (version 1.0.7) (Höhna et al. 2016) for each independent locus following the developers’ guidelines (Höhna et al. 2017). We applied a general time reversible (GTR) substitution model (Tavaré 1986) with flat Dirichlet prior distributions on both the stationary frequencies and on the exchangeability rates (Abadi et al. 2019). We modelled among-site rate variation using a discretized gamma distribution with four rate categories (Yang 1994) and with a lognormal prior distribution on the parameter alpha with mean 2.0 and standard deviation 0.587405, see (Höhna et al. 2017). To estimate divergence times, we used a birth-death process as the prior distribution on the tree topology and divergence times (Yang and Rannala 1997). Furthermore, to account for rate variation among lineages we used a relaxed-clock model with uncorrelated lognormal distributed rate categories (UCLN) (Drummond et al. 2006). We applied a hyperprior distribution on both the mean ∼ lognormal (median=0.01, sd=0. 587405) and standard deviation, sd ∼ exponential (10), of the branch-specific clock rates.

A firefly fossil, dated from Dominican amber at 20-25 MYA, was used for the node calibration at the *P. pyralis* bifurcation (Poinar and Poinar 1999; Grimaldi and Engel 2005). Hence, we calibrated the root node of the clade using a normal distribution with mean 22.5 and standard deviation of two to account for the uncertainty of this calibration (Parham et al. 2012). We ran a four-replicate Markov chain Monte Carlo (MCMC) analysis, as implemented in RevBayes, for 50,000 iterations and sampled phylogenetic trees and parameters states every 10 iterations, yielding a total of 20,000 samples from the posterior distribution. To ensure convergence of the replicated MCMC analyses, we made sure every parameter had an effective sample size (ESS) of at least 200 using the software *Tracer* (version 1.7) (Rambaut et al. 2018).

## Results

### Population clustering according to genetic similarity

We used published SNP data generated for 12 populations from the US, yielding a total of 308 sequenced chromosomes, to elucidate the demographic history of *P. pyralis* (Figure 1A) (Lower et al. 2018). Population-structure and genetic-distance analysis (Neighbor-joining) done in Lower et. al. 2018 suggested that the sampled populations can be clustered in three main population groups: the Texan cluster (including two Texan populations), a Central (Arkansas, Mississippi, Missouri, Ohio and Michigan), and an Eastern cluster (Georgia, Tennessee, North Carolina and New Jersey) (Table 1). To examine whether these three clusters can be correctly assigned by their genetic relatedness using all sampled populations and to statistically evaluate for the population grouping as assigned in Lower et al. 2018, we used a <u>D</u>iscriminant <u>A</u>nalysis of <u>P</u>rincipal <u>C</u>omponents (DAPC) (Jombart et al. 2010). Our DAPC analysis supported the three above mentioned clusters (Figure 1B) with high posterior probabilities for all samples (Table S1). Therefore, we used these three population clusters for the demographic inference. Throughout the rest of the paper we will refer these as the Texan (grey), the Central (blue) and the Eastern (light blue) populations (Figure 1).

**Figure 1.**
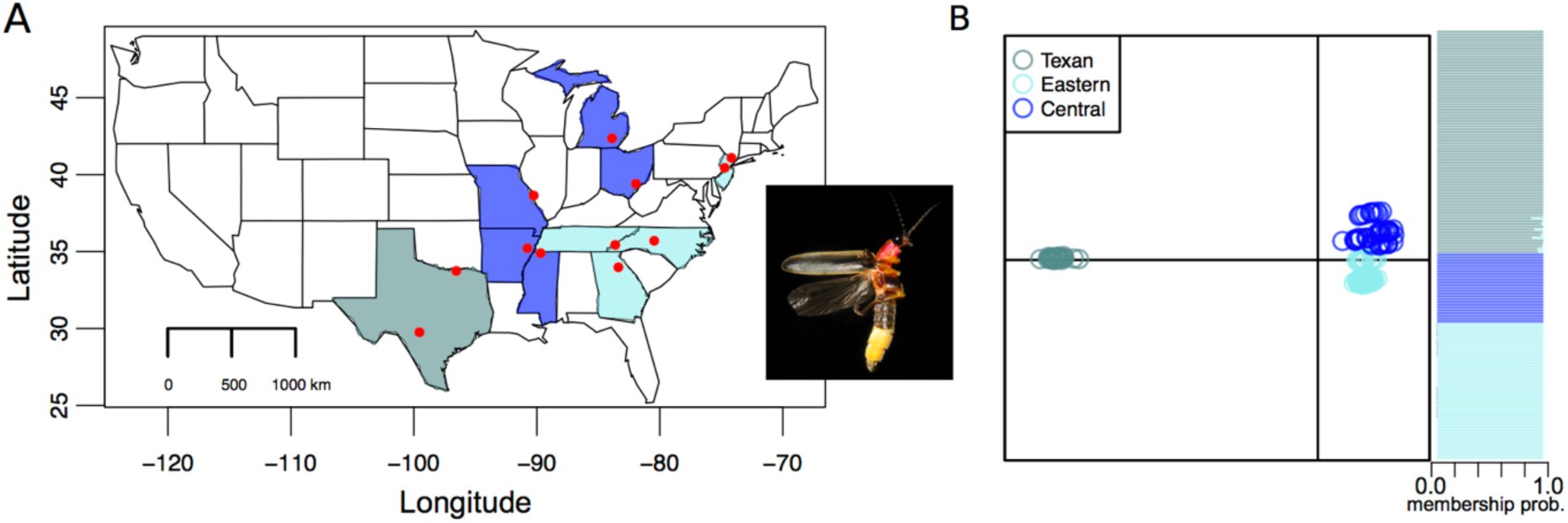
(A) Map of the US showing with red dots the sampled populations of *P. pyralis* (Lower et al. 2018). Y-axis shows the latitude (N°) and the x-axis shows the longitude (W°). The three colors represent the three population clusters used in this study. Grey: Texan population, blue: Central and light blue: Eastern population. Inset: *Photinus pyralis*, credit: Creative Commons. (B) Genetic clustering of all sampled individuals into three distinct genetic clusters defined by a Discriminant Analysis of Principal Components (DAPC), including cluster probabilities. Each dot represents a single individual (Texas: N = 25, Central = 49, Eastern = 80; Total: 153).

### Population genetics summary statistics

We calculated the nucleotide diversity estimators *θ*_W_ and *π* from our filtered data and observed that the Texan population has the highest nucleotide diversity when compared to the Central and Eastern populations (Table 2). The Texan population has 1.4- and 1.9-fold higher average number of pairwise differences (as estimated by *π*) than the Central and the Eastern population, respectively. When looking at the nucleotide diversity as estimated by the number of segregating sites, *θ*_W,_ the Texan population showed 1.05- and 1.21-fold higher values than the Central and the Eastern populations, respectively. Additionally, when looking at linkage disequilibrium levels as measured by *Z*_nS_ the Texan and the Eastern populations have comparable levels of linkage, which contrasted with the lower levels of linkage found in the Central population (Table 2). When calculating Tajima’s *D*, the Texan population has a value that is closest to zero, whereas the Central and the Eastern populations have a strong negative Tajima’s *D* (Table 2).

Next, we calculated the Wakeley-Hey W statistics in order to estimate the degree of isolation between populations by summarizing the joint site frequency spectrum (JSFS). From this analysis, when looking at private and shared polymorphisms between populations (W1, W2, W4), we found that the Central and Eastern populations are the closest to one another, when compared to the Texan population (Table 3). We also observed that there are almost no fixed differences between any pair of populations, and that most of the polymorphisms present in this dataset are either private or shared between each pair of populations (Table 3). In agreement with the genetic distance analysis done in Lower et. al. 2018 (Neighbor-joining), our estimation of the genetic distance (distance of Nei, Nei and Li 1979) also shows that the Texan population has the highest degree of genetic differentiation in comparison to the Central and the Eastern populations (Table 3).

Taken together that: (1) the Texan population has the highest nucleotide diversity levels and (2) the Texan population shows the most genetic differentiation from the rest of the populations we hypothesize that the Texan population is more ancestral with regard to the Central and Eastern populations.

### Demographic models

To test whether the Texan population is the most ancestral population in our North American firefly dataset we tested for four demographic scenarios using ABC: *Model 1:* The Texan population as the ancestral population, with subsequent sequential colonization of the Central and Eastern populations. *Model 2*: The Eastern population as the ancestral population, with subsequent sequential colonization of the Central and Texan populations. *Model 3*: The Texan as the ancestral population, with independent colonization of the Central and Eastern populations. *Model 4*: The Central population as the ancestral population, with independent colonization of the Texan and Eastern populations (Figure 2). With these different demographic scenarios we covered all biologically plausible population histories of North American *P. pyralis* based on population structure, known phylogeographic breaks and patterns of mitochondria COI data (Lower et al. 2018).

**Figure 2.**
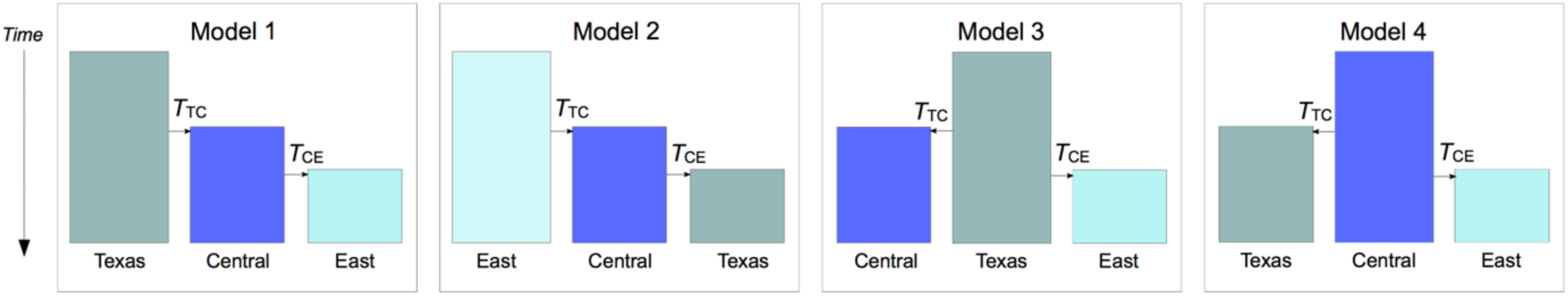
Demographic history models tested for *P. pyralis*. Boxes represent effective populations sizes (*Ne*) of each population. *T*_CE_ corresponds to the split time between the Central and Eastern populations, *T*_TC_ corresponds to the split time between the Texan and Central populations. All population sizes are estimated with respect to *Ne*_Texas_, therefore *Ne*_Texas_=1.

### Model choice

Validation of model choice using simulated datasets shows a high power for selecting the right model among the four tested scenarios. More specifically, Models 1 through 4 are chosen correctly 94%, 87%, 91%, and 95% of the time, respectively. With this high validation power, we then proceeded to calculate the posterior probabilities of all four models given the observed data. We found that Model 1 (southern origin + sequential colonization) has the highest probability (99.9%), and to predict well all tested summary statistics, that is, in all cases the observed summary statistic fell within the distribution of simulated statistics under Model 1 (Figure S2)

### Parameter estimation

By using <u>p</u>artial least <u>s</u>quares (pls) we reduced the dimensionality of all summary statistics down to ten pls components. This procedure is favorable since the new set of transformed statistics are orthogonal to each other, guaranteeing the assumption of singularity, which is required for ABC regression (Beaumont et al. 2002). Validation of parameter estimation showed good power to estimate all parameters except for the selection coefficient S_Aa_ (Figure S1). However, provided there is good power to estimate the other parameters (population sizes and timing of population splits) we report these estimates below (Table 4). All estimates (except for SAa) are given relative to *Ne*_Texas_.

Overall, the data reflects that the Central population has an effective population size similar to that of the ancestral Texan population, and that the Eastern population is much smaller than the Texan population. The split between the Eastern and Central populations happened around 0.0048*4Ne_Texas_ generations ago (Table 4, Figure 3). Currently, we cannot estimate the size of the Texan population because we don’t know the mutation rate (recall that θ=4Ne***µ***, where ***µ*** is the mutation rate). Nevertheless, for illustration purposes, if we were to assume that this population contains around 100,000 individuals and that the generation time in *P. pyralis* is about 2 years, then the split between the Central and the Eastern populations would have happened around 4,000 years ago. The split between the Texan and Central populations took place around 0.026*4Ne_Texas_ generations ago (again, applying the same illustrative example concerning population size and generation time we would conclude that this split took place around 21,000 years ago). We cannot say much about the strength of selection on this data set since we have no power to estimate this parameter. Finally, alternative models with migration between populations have also been analyzed but there was not enough power to tell migration from no-migration models (Table S1).

**Figure 3.**
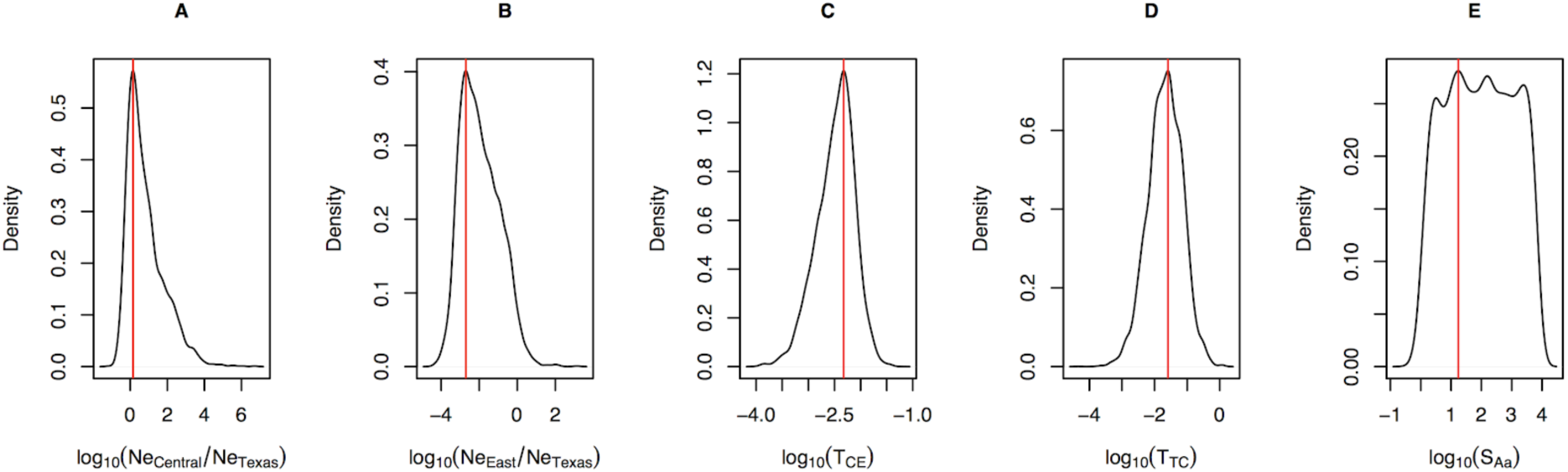
Posterior distribution of parameter estimates of Model 1. An explanation of the parameters and the characteristics of each distribution is detailed in Table 4.

### Demographic analysis for single population

Next, we investigated the putative demographic processes that each of the individual populations went through in their recent past. We used *dadi* (Gutenkunst et al. 2009) to test for five demographic scenarios (see Methods). The demographic model that had the lowest likelihood for our three populations (Texan, Central, and Eastern) was the constant population size scenario, clearly indicating that our firefly populations have gone through population changes. For all three populations the demographic models depicting population size changes had a very close fit (Table 5, Figure S3).

**Table 5.** Log-likelihoods of five demographic scenarios tested by *dadi*. For each population, bold indicates the demographic model with the highest likelihood.

| Demographic models | Texan population | Central population | Eastern population |
| --- | --- | --- | --- |
| (1) Neutral equilibrium | -340.62 | -540.38 | -375.94 |
| (2) Two-epoch | **-99.69** | -207.60 | -179.76 |
| (3) Growth | -99.84 | -208.50 | -180.00 |
| (4) Bottle growth | -99.76 | -237.25 | -180.00 |
| (5) Three epoch | -106.76 | **-205.12** | **-179.61** |

The best fitting model for the Texan population was the two-epoch model, suggesting that the Texan population underwent a population shrinkage 0.00026 2*N_e_* generations ago (Table 6). For the Central population the three-epoch model had the best fit. In this case, the Central population went first through a population expansion and then through a population shrinkage. The three-epoch model was also the best fit for the Eastern population, depicting a population expansion followed by a population decline (Table 6).

**Table 6.** Demographic parameter estimates for each population under the best fitted model.

| Population | Best model | $n_B^1$ | $n_F^2$ | $T_B^3$ | $T_F^4$ |
| --- | --- | --- | --- | --- | --- |
| Texan | Two epoch |  | 0.0001 |  | 0.00026 |
| Central | Three epoch | 28.9 | 9.5e-03 | 8.25 | 4.51e-03 |
| Eastern | Three epoch | 28.45 | 0.30 | 10.16 | 0.17 |

<sup>1</sup> Ratio of population size of the second epoch to the ancestral population.
<sup>2</sup> Ratio of population size of the third epoch to the ancestral population.
<sup>3</sup> Duration of the second epoch \* $2N_e u$ generations
<sup>4</sup> Duration of the third epoch \* $2N_e u$ generations

### Age of the most ancestral population of P. pyralis

To increase our understanding on the population history of *P. pyralis* we estimated the age of the phylogenetic node leading to *P. pyralis.* The age of the node to *P. pyralis* will give us an estimate of the age of its most ancestral population, possibly the Texan population or another population located more south of its distribution. For the estimation of the node age, we downloaded from GenBank all sequences with the taxon tag Lampyridae (fireflies) and built orthoclusters to identify loci to be used to generate a calibrated tree. We identified five such loci, three mitochondrial and two nuclear: *Cytochrome c oxidase subunit I* (COI, length:1478nt, sequences:123), *28S ribosomal RNA* (28S, length:632, sequences:28), *16S ribosomal RNA* (16S, length:473, sequences: 100), *wingless* (WG, length:404, sequences:55) and *carbamoyl-phosphate synthetase 2* (CAD, length:1763, sequences:41). By using a relaxed clock for the calibration and a firefly Dominican 20-25 MY-old fossil (Poinar and Poinar 1999; Grimaldi and Engel 2005) we were able to estimate the node age leading to *P. pyralis* for each of the five loci (Figure 5).

**Figure 5.**
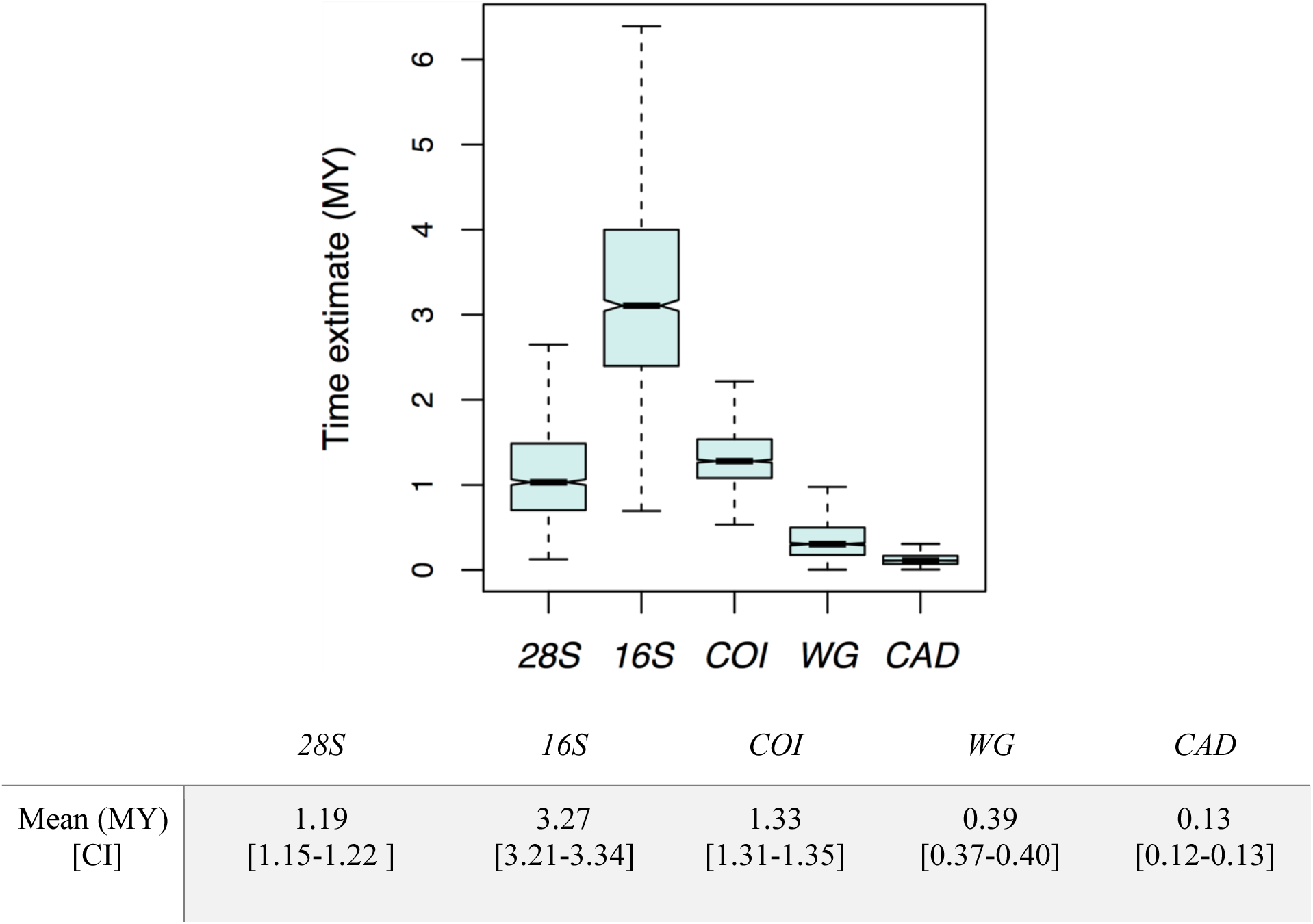
Boxplots show the age estimation in million years (MY) for the node leading to *P. pyralis* using a relaxed-clock model. Estimations are shown for three mitochondrial loci (*COI*, *16S* and *28S*) and two nuclear loci (*WG* and *CAD*). Boxplot areas show the first and third quartile of the estimated age values distribution and black bars indicate the median of the distribution. Whiskers indicate the lower and upper quartiles extending 1.5x the interquartile range.

The locus with the lowest node estimate is *CAD* with 0.1 million years (MY) and the one with the highest estimate is *16S* with 3.2 MY. The mitochondrial loci show a 5.1-fold higher age estimate when compared with the nuclear loci (mitochondrial loci age mean: 2.07 MY, nuclear loci age mean: 0.4 MY, Figure 5). As a result, we took the mean age of the node leading to *P. pyralis* from the five analyzed loci, which is ∼ 1.4 MY, as a proxy for the age of its most ancestral population.

## Discussion

### Demographic Inference of P. pyralis

The firefly P. pyralis has a wide distribution, which ranges from Ontario, Canada to South America (Venezuela, Brazil) (Fallon et al. 2018, The Bavarian State Zoological Collection). How did P. pyralis achieve this distribution pattern and where does the most ancestral population of the species lie? To start discerning the demographic history of P. pyralis we analyzed 12 populations sampled in North America (Figure 1A). From the four demographic hypothesis that we tested (Figure 2), Model 1 had the highest probability when compared with the rest of the models. Model 1 describes a scenario where the Texan population is the most ancestral population. The Central population is derived from the Texan one and in turn, the Eastern population is derived from the Central population. This stepwise colonization process, as depicted in Model 1, showed a 99.9% probability when compared to the 3 remaining models. Our study confidently rejects a demographic scenario in which colonization happened the other way around, from the Eastern US to Texas (Model 2). The demographic scenarios where the Central and the Eastern populations were independently colonized from Texas (Model 3) or where the Central population is the most ancestral one (Model 4), had zero model support. Placing Texas as the most ancestral population is also supported by the nucleotide diversity estimators θ_W_ and π, where Texas has the highest nucleotide diversity, followed by the Central population, and finally the Eastern population, with the lowest diversity values (Table 2).

The levels of LD vary between populations, and in this case, the Texan population has the highest LD, followed by the Eastern population, and finally the Central population with the lowest LD (Table 2). The strength of LD can increase due to gene flow between populations, (especially when allele frequencies differ among populations), when recombination rates are low or after a recent migration event (Slatkin 2008). Changes in population size can also influence LD levels, such as bottlenecks which could potentially increase LD levels due to the loss of alleles (Zhang et al. 2004). Finally, positive selection can also increase LD due to genetic hitchhiking during a selective sweep (Kim and Stephan 2002).

When looking at the Wakeley-Hey W statistics we observe a gradient in terms of shared polymorphisms, where the number of shared polymorphisms decreases with geographic distance (Table 3), an expected outcome in a demographic scenario were a stepwise colonization has taken place. Additionally, the fact that 32-43% of the SNPs are shared across populations, the low number of fixed differences between all populations and the low global F_st_ value, suggest that the colonization of the Central and the Eastern populations are recent events. Alternatively, high levels of shared polymorphisms and low F_st_ values across populations can also be obtained under the presence of gene flow (Wakeley and Hey 1997). The wide distribution of P. pyralis already suggests its high dispersion capacity, thus making the presence of gene flow between populations plausible. Consequently, we also tested models including migration but with our current RAD data, there was not enough power to distinguish between migration from no-migration models (Table S1). Migration models would benefit significantly by obtaining more SNPs and haplotypes sampled from whole genomes.

The population-specific statistics, Tajima’s D, LD, and the joint SFS, already suggest that each population might have undergone size changes. Consequently, by using dadi, we found that a constant size scenario had the lowest likelihood in our three populations. In the Central and the Eastern population the three-epoch model fitted best, showing that these populations went through expansions followed by shrinkage (Table 6). These results suggest that after the Central and the Eastern colonization, these populations went through a founder event that was followed by a recovery in population size, which is here depicted by a population expansion. This single population demographic scenario complements well with our ABC analysis, where the best model describes a stepwise colonization events of the Central and the Eastern population. From the three-epoch model, we can also see that after experiencing a population growth, both populations experienced a population shrinkage more recent in time. Single population demographic events can be very dynamic in nature, especially with constant climate changes and changes in habitat availability. As natural habitats usually have a tendency to shrink due to human impact, it is not entirely surprising that we found a signal of population shrinkage in the Central and in the Eastern populations. In the case of the Texan population, the two-epoch model fitted best, describing a decline in population size (Table 6). From our ABC demographic analysis we uncovered that the Texan population is the most ancestral P. pyralis population from North America. Being this population older than the Central and the Texan, the two-epoch model suggests that the Texan population had fully recovered from a founder event, as we do not observe a signal for a population expansion as in the Central and Eastern populations. The more recent population shrinkage that we observe in the Texan population could be linked to more recent events resulting in a population decline. The single population demographic inference using dadi has generated hypotheses on the putative demographic changes experienced by these populations. Nevertheless, given the close fit of all the models tested (Table 5), the above proposed demographic scenarios will have to be further tested when more data is available.

From our demographic modelling, the size of the Central population is slightly larger (∼1.43 times larger) than that of the Texan population. The Central population has the highest number of segregating sites and the lowest values of LD when compared to the other populations (Table 2). This is further evidence for a scenario where the Central population has experienced a population expansion as suggested in our dadi analysis. In the case of the Eastern population, it shows a population size 0.0020 smaller than that of the Texan population. The relatively small Eastern population size might be a result of a recent colonization event, putatively a post-glaciation event, or higher selective pressures that do not allow for bigger population sizes (e.g. winter). Equally, as shown by our single population demographic analysis, the small size of the Eastern population might be a result of a two-step population size change, where the Eastern population recovered from a putative founder event, stated by a population expansion in the three epoch model, followed by a population shrinkage (Table 6). The low and homogeneous genetic diversity of the Easter population (Table 2, Table 3, Lower et al. 2018) could also be maintained by the Appalachian mountain ranges serving as a geographic barrier between the Eastern population and the Central and Texan population (Soltis et al. 2006).

Furthermore, we estimated the relative time of colonization of the Central and Eastern populations in relation to the Texan effective population size. Nevertheless, because we do not have the actual estimate of the Texan population size, we can only express time estimates relative to the effective population size Ne of the Texan population. The Ne in other insects, for example Drosophila melanogaster (Arguello et al. 2019) and Heliconius melpomene (Keightley et al. 2015), has been estimated to be around 2 million individuals for both species. If the Ne of P. pyralis is close to 2 million individuals then we can estimate that the Central population was founded ∼ 400,000 years ago, and the Eastern ∼ 75,000 years ago. When using the lady beetle, Hippodamia convergens, as an example for Ne in beetles, the biggest population studied had a Ne of ∼11,000 individuals. If the Ne of P. pyralis would be in that range, the Central population would have been then colonized ∼3,000 years ago and the Eastern population only ∼500 years ago. Thus, the specific time of colonization of the Central and the Eastern population still remains to be elucidated in order to have more precise colonization times. Two variables will help to get a more accurate estimate of the colonization time. The first one will be estimating the mutation rate of P. pyralis, which is not a trivial task, but plausible by either generating pedigree lines or by generating population-level whole-genome data from which we can draw mutation rate estimates by using synonymous sites (Yang and Nielsen 1998), microsatellites (Whittaker et al. 2003) or pseudogene variation (Nachman and Crowell 2000). The second factor influencing the estimation of the time of colonization is the generation time of P. pyralis which is hypothesized to vary across populations. For example, the populations inhabiting northern latitudes such as the Eastern population only produce one generation every two years, whereas southern populations are hypothesized to produce two generations a year (Faust 2017; Fallon et al. 2018). More research on the life cycle of P. pyralis will help to get more precise values of the generation time, which we can then include in our demographic models.

The estimation of the age of P. pyralis’s most ancestral population gives us an anchor in time of the evolutionary history of the species. From nuclear and mitochondrial loci (Figure 5) we have estimated that the age of the node to P. pyralis ranges from ∼0.125 to 3.3 million years, depending on the locus tested. The three mitochondrial loci that we used resulted in an older node age (1.2 to 3.3 million years) as opposed to the estimates recovered from the nuclear loci (∼0.125,189 - 386,274 years). This discrepancy might be explained by the mitochondria having higher mutation rates than nuclear genes (Moriyama and Powell 1997) and overall by the differences in evolutionary change within a genome (Hodgkinson and Eyre-Walker 2011). Between 125,000 and 3.3 million years ago the land bridge of Central America had been completed hinting that P. pyralis’s dispersion from South to North America was not hindered by big stretches of sea (Iturralde-Vinent 2006). From these results we hypothesize that the Pleistocene glaciation (∼2.58 MYA to the present) had the most impact on P. pyralis current geographical distribution, specially the glaciation events occurring during the last glacial period (115,000 – 11,700 years ago) (Eglinton et al. 2005). Some of the locations where our populations were collected (e.g., New Jersey and Michigan), were covered by ice during the last glaciation (Bemmels and Dick 2018), making these areas habitable only after a glacial retreat. The future estimation of effective population sizes in P. pyralis will help us to more accurately infer the colonization times of the Central and Eastern populations as well as to better understand the role of glacial refugia in P. pyralis’ current distribution.

### Alternative demographic scenarios

Our study on the demographic history of *P. pyralis* supports the hypothesis that this species has a southern origin and that, recently colonized the central and the north eastern part of North America in a stepwise manner. However, *P. pyralis* distribution goes further south, going through Mexico and Central America down to Venezuela and Brazil (Fallon et al. 2018, The Bavarian State Zoological Collection). Therefore, as samples from Central and South America become available, we will be able to test at least two more demographic scenarios: (1) Speciation of *P. pyralis* taking place in Nuclear Central America followed by independent migration events to North and South America; (2) the most ancestral population originating in South America followed by gradual migration to the North. Nuclear Central America is a particularly speciose area, with high endemism where *in situ* diversification of supraspecific taxa has taken place (Cano et al. 2018), making this a candidate region for firefly diversification. On the other hand, a fair amount of biota diversified in South America and after the closure of the Panamá Isthmus, the great American biota interchange occurred in many waves, one of which *P. pyralis* could have taken to reach northern latitudes (Cione et al. 2015). Our present and future work on the demographic history of *P. pyralis* will enlighten us on the complex history of this species, its place of origin, past migration events and present population dynamics. Additionally, our work has established neutral expectations of genetic diversity for the North American populations, which can be set as a null model to investigate natural selection events.

## Author contributions

AC and SH conducted phylogenetic and *dadi* analysis. AC wrote the initial and final draft. PD performed demographic analyses and writing. SL contributed data, demographic models, and writing. All authors conceived the study and approved the final version of the manuscript.

## Acknowledgments

Jochen Wolf for supporting the project through the Knut & Alice Wallenberg Foundation granted to him. This research was additionally supported by the Deutsche Forschungsgemeinschaft (DFG) Emmy Noether-Program HO6201/1-1 awarded to SH.

## Data Accessibility Statement

archival location upon acceptance or statement that there is no data to be archived.

## Supplementary materials

**Figure S1.**
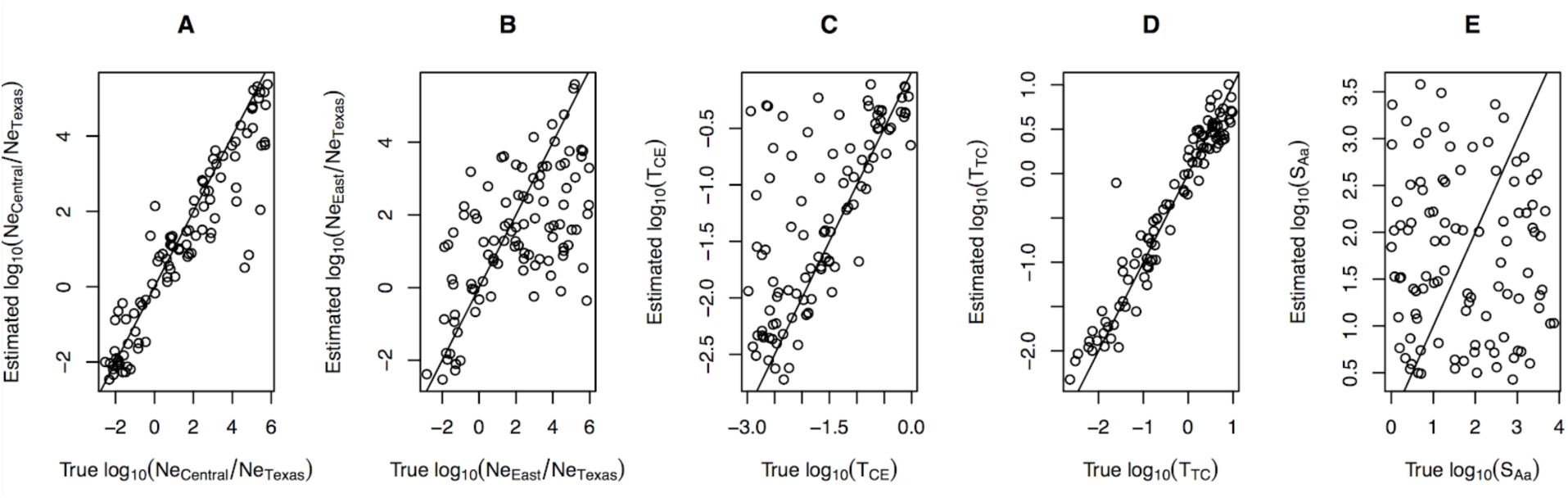
Validation of parameter estimation under Model 1. Here, true parameter values drawn from the simulations are plotted along the x-axis and the estimated parameters under ABC-regression are plotted in the y-axis. A positive correlation along a line with slope 1 (solid line) is expected if there is good power to estimate a parameter.

**Figure S2.**
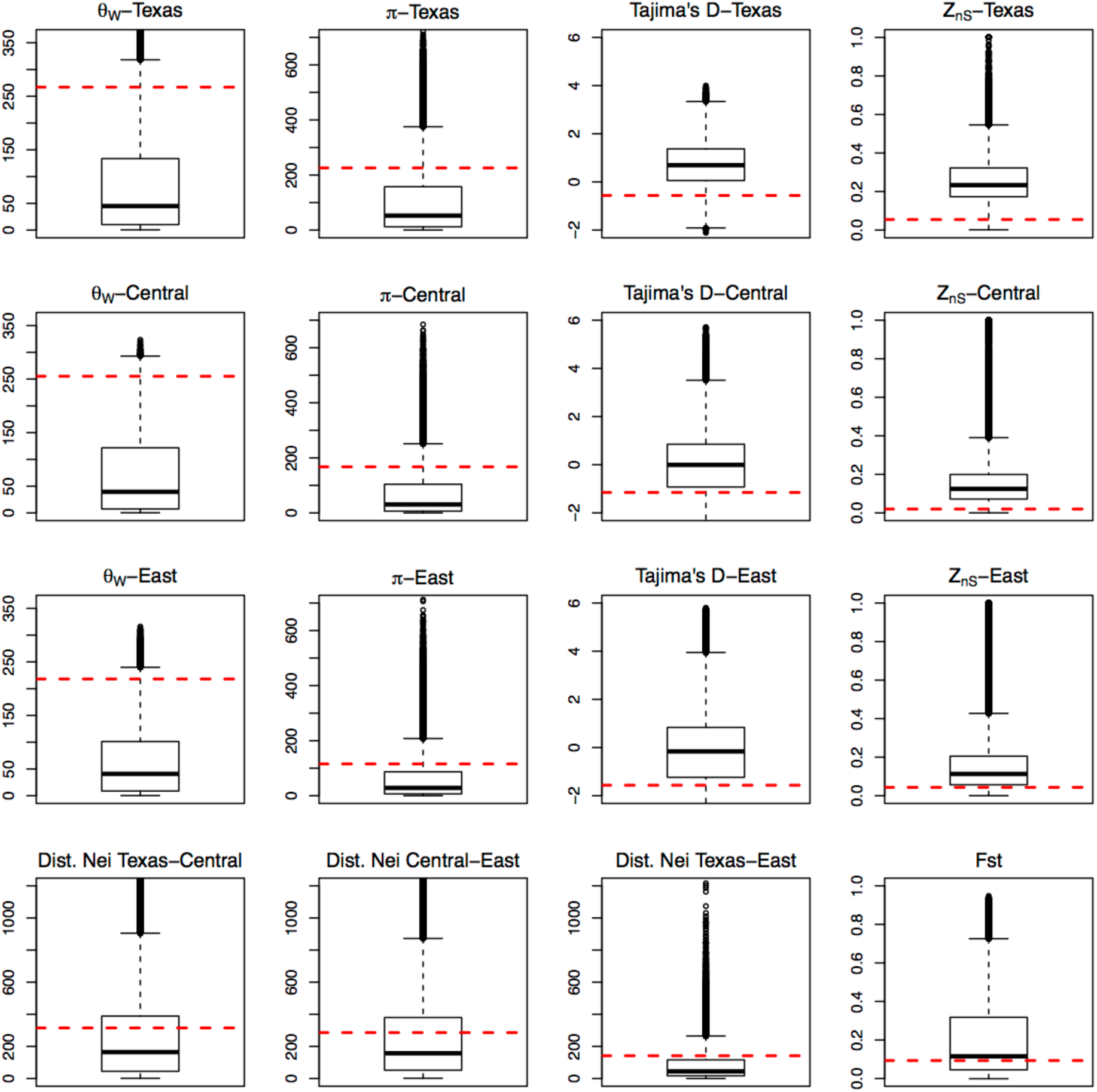
Predictive simulations for some key summary statistics. Each boxplot is based on the simulated summary statistics under Model 1. The corresponding observed statistic is shown with a red horizontal dashed line.

**Figure S3.**
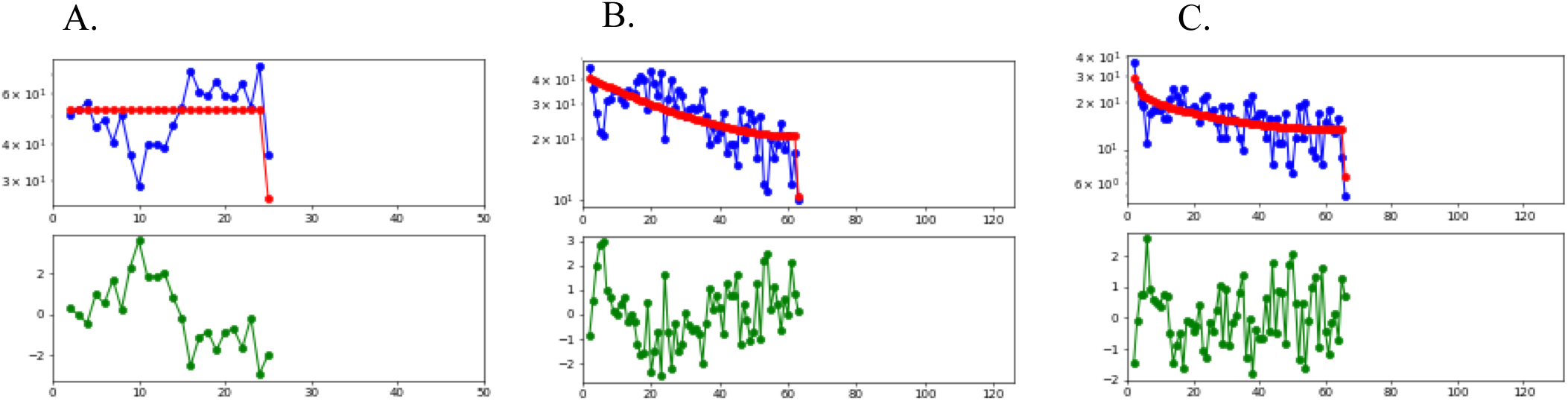
Upper panel: Calculation of the expected site frequency spectrum as calculated in *dadi* (red); calculated site frequency spectrum of three *P. pyralis* populaitons (blue). Y-axis shows the frequency of the site polymorphism clasees, x-axis shows the polymorphism classes drawn from a folded site frequency spectrum. Lower panel: In green is shown the residuals drawn from the correlation between the expected and the observed site frequency spectrum. A. Texan population: 2-epoch, B. Central population: 3-epoch, C. Eastern population: 3-epoch.

**Table S1.** Confusion matrix for the validation of model choice between migration and non-migration models. Model S1: Sequential colonization Texas-Central-Eastern; Model S2: same as S1 plus migration between Texas-Central and Central-Eastern; Model S3: Independent colonization Texas-Central and Texas-Eastern; Model S4: same as S3 plus migration between Texas-Central and Central-Eastern. As shown in this table, there is power to tell between the colonization scenarios, but not between migration vs no-migration.

|  | Model S1 | Model S2 | Model S3 | Model S4 |
| --- | --- | --- | --- | --- |
| Model S1 | 60 | 37 | 2 | 1 |
| Model S2 | 52 | 47 | 0 | 1 |
| Model S3 | 2 | 6 | 56 | 36 |
| Model S4 | 5 | 6 | 44 | 45 |

## Literature cited

Abadi, S., D. Azouri, T. Pupko, and I. Mayrose. 2019. Model selection may not be a mandatory step for phylogeny reconstruction. Nat. Commun. 10. Springer US.

Beaumont, M. A., W. Zhang, and D. J. Balding. 2002. Approximate Bayesian computation in population genetics. Genetics 162:2015–2035.

Bemmels, J. B., and C. W. Dick. 2018. Genomic evidence of a widespread southern distribution during the Last Glacial Maximum for two eastern North American hickory species. J. Biogeogr. 45:1739–1750.

Branham, M. A., and J. W. Wenzel. 2003. The origin of photic behavior and the evolution of sexual communication in fireflies (Coleoptera: Lampyridae). Cladistics 19:1–22.

Cano, E. B., J. C. Schuster, and J. J. Morrone. 2018. Phylogenetics of *Ogyges kaup* and the biogeography of nuclear central America (Coleoptera, Passalidae). Zookeys 2018:81–111.

Cione, A. L., G. M. Gasparini, E. Soibelzon, L. H. Soibelzon, and E. P. Tonni. 2015. The Great American Biotic Interchange: A South American Perspective. First edit. Springer US, London.

Csilléry, K., O. François, and M. G. B. Blum. 2012. Abc: An R package for approximate Bayesian computation (ABC). Methods Ecol. Evol. 3:475–479.

Drummond, A. J., S. Y. W. Ho, M. J. Phillips, and A. Rambaut. 2006. Relaxed phylogenetics and dating with confidence. PLoS Biol. 4:699–710.

Duchen, P., D. Zivkovic, S. Hutter, W. Stephan, and S. Laurent. 2013. Demographic inference reveals African and European admixture in the North American *Drosophila melanogaster* population. Genetics 193:291–301.

Edgar, R. C. 2010. Search and clustering orders of magnitude faster than BLAST. Bioinformatics 26:2460–2461.

Eglinton, G., A. Rosell-Mele, R. Tiedemann, M. A. Maslin, D. M. Sigman, M. J. Leng, S. L. Jaccard, G. E. A. Swann, J. Bollmann, G. H. Haug, and A. Ganopolski. 2005. North Pacific seasonality and the glaciation of North America 2.7 million years ago. Nature 433:821–825.

Ewing, G., and J. Hermisson. 2010. MSMS: A coalescent simulation program including recombination, demographic structure and selection at a single locus. Bioinformatics 26:2064–2065.

Excoffier, L., I. Dupanloup, E. Huerta-Sánchez, V. C. Sousa, and M. Foll. 2013. Robust Demographic Inference from Genomic and SNP Data. PLoS Genet. 9:e1003905. Public Library of Science (PLoS).

Fallon, T. R., S. E. Lower, C.-H. Chang, M. Bessho-Uehara, G. J. Martin, A. J. Bewick, M. Behringer, H. J. Debat, I. Wong, J. C. Day, A. Suvorov, C. J. Silva, K. F. Stanger-Hall, D. W. Hall, R. J. Schmitz, D. R. Nelson, S. M. Lewis, S. Shigenobu, S. M. Bybee, A. M. Larracuente, Y. Oba, and J.-K. Weng. 2018a. Firefly genomes illuminate parallel origins of bioluminescence in beetles. Elife 7:1–146.

Fallon, T. R., S. E. Lower, C. Chang, M. Bessho-Uehara, G. J. Martin, A. J. Bewick, M. Behringer, H. J. Debat, I. Wong, J. C. Day, A. Suvorov, C. J. Silva, K. F. Stanger-Hall, D. W. Hall, R. J. Schmitz, D. R. Nelson, S. M. Lewis, S. Shigenobu, S. M. Bybee, A. M. Larracuente, Y. Oba, J.-K. Weng, and J. Robert. 2018b. Firefly genomes illuminate the origin and evolution of bioluminescence. Elife 7:1–146.

Faust, L. F. 2017. Fireflies, glow-worms, and lightning bugs : identification and natural history of the fireflies of the eastern and central United States and Canada.

Freyman, W. A. 2015. SUMAC: Constructing phylogenetic supermatrices and assessing partially decisive taxon coverage. Evol. Bioinforma. 11:263–266.

Gonzalo, H. 1987. Biogeography of the montane entomofauna of Mexico and Central America. Annu. Rev. Entomol. 32:95–114.

Grimaldi, D., and M. S. Engel. 2005. Evolution of the insects. First edit. Cambridge University Press, Hong Kong.

Gutenkunst, R. N., R. D. Hernandez, S. H. Williamson, and C. D. Bustamante. 2009. Inferring the Joint Demographic History of Multiple Populations from Multidimensional SNP Frequency Data. PLoS Genet 5:e1000695. Public Library of Science.

Hodgkinson, A., and A. Eyre-Walker. 2011. Variation in the mutation rate across mammalian genomes. Nat. Rev. Genet. 12:756–766. Nature Publishing Group.

Höhna, S., M. Landis, and H. T. A. 2017. Phylogenetic Inference Using RevBayes. Curr. Protoc. Bioinforma. 57:6.16.1–6.16.34. Wiley-Blackwell.

Höhna, S., M. J. Landis, T. A. Heath, B. Boussau, N. Lartillot, B. R. Moore, J. P. Huelsenbeck, and F. Ronquist. 2016. RevBayes: Bayesian phylogenetic inference using graphical models and an interactive model-specification language. Syst. Biol. 65:726–736.

Iturralde-Vinent, M. A. 2006. Meso-Cenozoic Caribbean Paleogeography: Implications for the Historical Biogeography of the Region. Int. Geol. Rev. 48:791–827.

Jombart, T., and I. I. Ahmed. 2011. adegenet 1.3-1: new tools for the analysis of genome-wide SNP data. Bioinformatics 27:3070–3071.

Jombart, T., S. Devillard, and F. Balloux. 2010. Discriminant analysis of principal components: a new methods for the analysis of genetically structured populations. BMC Genet. 11:1–15.

Katoh, K., K. Misawa, K. Kuma, and T. Miyata. 2002. MAFFT: a novel method for rapid multiple sequence alignment based on fast Fourier transform. Nucleic Acids Res. 30:3059– 3066.

Keightley, P. D., A. Pinharanda, R. W. Ness, F. Simpson, K. K. Dasmahapatra, J. Mallet, J. W. Davey, and C. D. Jiggins. 2015. Estimation of the spontaneous mutation rate in *Heliconius melpomene*. Mol. Biol. Evol. 32:239–243.

Kelly, J. K. 1997. A test of neutrality based on interlocus associations. Genetics 146:1197–206.

Kim, Y., and W. Stephan. 2002. Detecting a local signature of genetic hitchhiking along a recombining chromosome. Genetics 160:765–77.

Larsson, A. 2014. AliView: A fast and lightweight alignment viewer and editor for large datasets. Bioinformatics 30:3276–3278.

Lessios, H. A. 2008. The great American schism: Divergence of marine organisms after the rise of the Central American Isthmus. Annu. Rev. Ecol. Evol. Syst. 39:63–91.

Lewis, S. M., and C. K. Cratsley. 2008. Flash Signal Evolution, Mate Choice, and Predation in Fireflies. Annu. Rev. Entomol. 53:293–321.

Lloyd, J. E. 1966. Studies on the flash communication system in Photinus fireflies. Misc. Publ. Museum Zool. Univ. Michigan 130:1–95.

Lower, S. E., K. F. Stanger-hall, and D. W. Hall. 2018. Molecular variation across populations of a widespread North American firefly, *Photinus pyralis*, reveals that coding changes do not underlie flash color variation or associated visual sensitivity. BMC Evol. Biol. 18:1–14. BMC Evolutionary Biology.

Mann, P. 2007. Overview of the tectonic history of northern CentralAmerica. Geol. Soc. Am. spec. pap. 428:1–19.

Martin, G. J., M. A. Branham, M. F. Whiting, and S. M. Bybee. 2017. Total evidence phylogeny and the evolution of adult bioluminescence in fireflies (Coleoptera: Lampyridae). Mol. Phylogenet. Evol. 107:564–575. Elsevier Inc.

Moriyama, E. N., and J. R. Powell. 1997. Synonymous substitution rates in *Drosophila*: Mitochondrial versus nuclear genes. J. Mol. Evol. 45:378–391.

Nachman, M. W., and S. L. Crowell. 2000. Estimate of the mutation rate per nucleotide in Humans. Genetics 156:297–304.

Nei, M., and W. H. Li. 1979. Mathematical model for studying genetic variation in terms of restriction endonucleases. Proc Natl Acad Sci U S A 76:5269–5273.

Parham, J. F., P. C. J. Donoghue, C. J. Bell, T. D. Calway, J. J. Head, P. A. Holroyd, J. G. Inoue, R. B. Irmis, W. G. Joyce, D. T. Ksepka, J. S. L. Patané, N. D. Smith, J. E. Tarver, M. Van Tuinen, Z. Yang, K. D. Angielczyk, J. M. Greenwood, C. A. Hipsley, L. Jacobs, P. J. Makovicky, J. Müller, K. T. Smith, J. M. Theodor, R. C. M. Warnock, and M. J. Benton. 2012. Best practices for justifying fossil calibrations. Syst. Biol. 61:346–359.

Poinar, G., and R. Poinar. 1999. The amber forest, a reconstruction of a vanished world. first. Princeton University Press, Princeton, NJ, NJ.

Pritchard, J. K., M. T. Seielstad, and M. W. Feldman. 1999. Population growth of human Y chromosomes: A study of Y chromosome microsatellites. Mol. Biol. Evol. 16:1791–1798.

Rambaut, A., A. J. Drummond, D. Xie, G. Baele, and M. A. Suchard. 2018. Posterior summarisation in Bayesian phylogenetics using Tracer 1.7. Syst. Biol. 00:1–3.

Roy, A. J., and M. S. Lachniet. 2010. Late Quaternary glaciation and equilibrium-line altitudes of the Mayan Ice Cap, Guatemala, Central America. Quat. Res. 74:1–7.

Sander, S. E., and D. W. Hall. 2015. Variation in opsin genes correlates with signalling ecology in North American fireflies. Mol. Ecol. 24:4679–4696.

Schuster, J. C. 1997. Seasonal diversity of fireflies (Coleoptera:Lampyridae) in a montane area of Guatemala. Proc. Int. Symp. Biodivers. Syst. Trop. Ecosyst. 281–284.

Slatkin, M. 2008. Linkage disequilibrium - Understanding the evolutionary past and mapping the medical future. Nat. Rev. Genet. 9:477–485.

Soltis, D. E., A. B. Morris, J. S. McLachlan, P. S. Manos, and P. S. Soltis. 2006. Comparative phylogeography of unglaciated eastern North America. Mol. Ecol. 15:4261–4293.

Stanger-Hall, K. F., and J. E. Lloyd. 2015. Flash signal evolution in Photinus fireflies: Character displacement and signal exploitation in a visual communication system. Evolution (N. Y). 69:666–682.

Stanger-Hall, K. F., J. E. Lloyd, and D. M. Hillis. 2007. Phylogeny of North American fireflies (Coleoptera: Lampyridae): Implications for the evolution of light signals. Mol. Phylogenet. Evol. 45:33–49.

Tajima, F. 1989. Statistical Method for Testing the Neutral Mutation Hypothesis by DNA Polymorphism. Genetics 595:585–595. Genetics Society of America.

Talavera, G., and J. Castresana. 2007. Improvement of phylogenies after removing divergent and ambiguously aligned blocks from protein sequence alignments. Syst. Biol. 56:564–577.

Tavaré, S. 1986. Some probabilistic and statistical problems in the analysis of DNA sequences. Am. Math. Soc. Lect. Math. Life Sci. 17:57–86.

Tavare, S., D. J. Balding, J. R. C. Griffiths, and P. Donnelly. 1997. Inferring Coalescence TimesFrom DNA Sequence. Genetics 145:505–518.

Vencl, F. V, and A. D. Carlson. 1998. Proximate mechanisms of sexual selection in the firefly *Photinus pyralis* (Coleoptera : Lampyridae). J. Insect Behav. 11:191–207.

Wakeley, J., and J. Hey. 1997. Estimating ancetral population parameters. Genetics 145:847–855.

Watterson, G. A. 1975. On the number of segregating sites in genetical models without recombination. Theor. Popul. Biol. 276:256–276.

Weir, B. S., and C. C. Cockerham. 1984. Estimating F-statistics for the analysis of population structure. Evolution (N. Y). 38:1358–1370.

Whittaker, J. C., R. M. Harbord, N. Boxall, I. Mackay, G. Dawson, and R. M. Sibly. 2003. Likelihood-based estimation of microsatellite mutation rates. Genetics 164:781–787.

Yang, Z. 1994. Maximum likelihood phylogenetic estimation from DNA sequences with variable rates over sites: Approximate methods. J. Mol. Evol. 39:306–314.

Yang, Z., and R. Nielsen. 1998. Synonymous and nonsynonymous rate variation in nuclear genes of mammals. J. Mol. Evol. 46:409–418.

Yang, Z., and B. Rannala. 1997. Monte Carlo Method A Markov Chain I-I. Integr. Vlsi J. 14:717–724.

Zhang, W., A. Collins, J. Gibson, W. J. Tapper, S. Hunt, P. Deloukas, D. R. Bentley, and N. E. Morton. 2004. Impact of population structure, effective bottleneck time, and allele frequency on linkage disequilibrium maps. Proc. Natl. Acad. Sci. 101:18075–18080.

